# Lysophosphatidic acid receptor 1 influences disease severity in a mouse model of multiple sclerosis

**DOI:** 10.1101/2024.05.28.596148

**Authors:** Nagi Uemura, Nagisa Matsuo, Kazunao Taniyama, Hiroki Nakajima, Kazuki Nagayasu, Shuji Kaneko, Jerold Chun, Hiroshi Ueda, Hisashi Shirakawa

## Abstract

Multiple sclerosis (MS), a chronic inflammatory disease affecting the central nervous system (CNS), is characterized by demyelination and axonal degeneration. Current treatments, which focus mainly on reducing lymphocyte infiltration into the CNS, are insufficient due to serious side effects and limited effectiveness; thus, identifying drugs with new mechanisms of action is crucial. Lysophosphatidic acid (LPA), a bioactive lipid produced by the enzyme autotaxin, may play a role in MS pathogenesis. Specifically, the LPA_1_ subtype of LPA receptors is linked to release of inflammatory cytokines in the CNS, and to demyelination in the peripheral nervous system. Our study investigated the role of LPA_1_ in a mouse model of MS. Knocking out the LPA_1_ gene in mice with experimental autoimmune encephalomyelitis improved clinical outcomes and reduced demyelination. Additionally, the absence of LPA_1_ reduced activation of Iba1-positive cells. Treatment with AM095, an LPA_1_ antagonist, tended to improve clinical outcomes and reduce levels of inflammatory mediators. These findings indicate that activation of LPA_1_ contributes to MS pathogenesis by promoting microglial activation and infiltration of peripheral immune cells.

## Introduction

Multiple sclerosis (MS), a chronic inflammatory, demyelinating, and neurodegenerative disease of the central nervous system (CNS) is believed to be autoimmune in nature [1,2]. Current therapies primarily target lymphocytes; however, these can lead to severe side effects and have proven inadequate for management of progressive MS [3,4]. Consequently, there is a pressing need for further investigation into the pathological mechanisms, as well as development of new treatments. The experimental autoimmune encephalomyelitis (EAE) mouse model is utilized extensively in MS research because it simulates many aspects of the disease [5,6]. Research indicates that myelin-specific autoimmune T helper (Th) cells, once activated in the periphery, migrate to the CNS where they trigger inflammation and subsequent demyelination [2,7]. Additionally, studies highlight the characteristic activation of monocyte lineage cells such as macrophages and microglia in the CNS of both patients with MS and EAE mice [8,9].

Lysophosphatidic acid (LPA) is a bioactive lipid synthesized predominantly from lysophosphatidylcholine by the enzyme autotaxin (ATX) [10]. Inhibition of ATX in EAE models improves pathology [11,12], suggesting that LPA biosynthesis and its receptor signaling pathways are key contributors to EAE pathogenesis. Previous studies report elevated levels of LPA in the serum and cerebrospinal fluid of MS patients, along with increased ATX activity in these fluids [13–15]. LPA interacts with at least six G protein-coupled receptors (LPA_1_–LPA_6_), thereby triggering various downstream signaling pathways [16,17]. Of these, LPA_1_ is expressed in multiple tissues, including the brain, thymus, and spleen, and it binds to three types of heterotrimeric G_α_ proteins: G_i/o_, G_q/11_, and G_12/13_ [18]. Inhibition of LPA_1_ reduces microglial activation and TNFα release in a systemic lipopolysaccharide (LPS)-induced septic model [19], and it also lessens demyelination of the spinal dorsal root in peripheral neuropathic pain models [20–23].

In this study, we investigated involvement of LPA_1_ in EAE, a mouse model of MS characterized by demyelination of the CNS, using genetically engineered mice and a pharmacological intervention.

## 2. Material and methods

### 2.1 Animals

All experiments were conducted in accordance with the ethical guidelines of the Kyoto University animal experimentation committee and the guidelines of the Japanese Pharmacological Society. The female C57BL/6J mice, LPA_1_ homozygous knockout (LPA_1_-homo-KO) mice, and LPA_1_ heterozygous knockout (LPA_1_-hetero-KO) mice (aged 7–12 weeks, weighing 18–22 g) [24] used in this study were maintained in specific-pathogen-free conditions. LPA_1_-homo-KO mice were backcrossed with C57BL/6J mice for at least ten generations to minimize any background effects on the phenotype. LPA_1_-homo-KO mice were then crossed with C57BL/6J mice to produce LPA_1_-hetero-KO mice. These LPA_1_-hetero-KO mice were bred with each other to generate LPA_1_-hetero-KO mice and their wild-type (WT) littermate controls. The mice were housed at a constant ambient temperature of 22 ± 2°C with a 12-h light/dark cycle, and had free access to food and water.

### 2.2 EAE

EAE was induced as described previously [6,25,26]. Briefly, mice were immunized subcutaneously with 100 µg of myelin oligodendrocyte glycoprotein peptide 35–55 (MOG 35-55; Scrum) in complete Freund’s adjuvant (231131, Difco) enriched with 3 mg/mL *Mycobacterium tuberculosis* H37Ra (231141, Difco). Additionally, mice received 10 µg/kg pertussis toxin (P7208, Sigma-Aldrich) intraperitoneally on the day of immunization, and again 48 h later. Clinical signs were scored daily using the following scale: 0, no clinical deficit; 1, partial tail paralysis; 2, full tail paralysis; 3, partial hind limb paralysis; 4, full hind limb paralysis; 5, forelimb paresis; 6, mortality.

The cumulative score was calculated as the sum of daily scores from Day 0 to Day 28.

### 2.3 FluoroMyelin staining

Mice were anesthetized with a mixture of 0.3 mg/kg medetomidine (Zenoaq), 4.0 mg/kg midazolam (Astellas Pharma), and 5.0 mg/kg butorphanol (Meiji-Seika Pharma). They were then perfused through the ascending aorta with phosphate-buffered saline (PBS), followed by 4% paraformaldehyde (PFA). Subsequently, lumbar spinal cords (L3–L5) were extracted and dissected. After post-fixation in 4% PFA for 3 h followed by cryoprotection overnight in 15% sucrose at 4°C, coronal sections (20 µm thickness) were prepared using a cryomicrotome (Leica). For FluoroMyelin staining, spinal cord sections were rinsed in PBS for 10 min, incubated with FluoroMyelin™ Green Fluorescent Myelin Stain (F34651, 1:1000; Invitrogen) for 20 min at room temperature, and then mounted with DAPI Fluoromount-G (Southern Biotech). Images were captured under a FluoView FV10i confocal microscope (Olympus). The areas of demyelination and total white matter were quantified using ImageJ software (National Institutes of Health). As described previously [26], percentage demyelination was calculated using the formula: demyelinated area (%) = (demyelinated area in white matter) / (total white matter area) × 100 (%).

### 2.4 Immunohistochemistry

Coronal sections were treated with 0.1% Triton X-100 in PBS for 15 min, followed by blocking in PBS containing 3% bovine serum albumin for 1 h. The sections were then incubated overnight at 4°C with the following primary antibodies at a dilution of 1:200: anti-Iba1 (019-19741, rabbit IgG; Wako Pure Chemical Industries), anti-CD3 (555273, rat IgG; BD Biosciences), and anti-Gr1 (MAB1037-100, rat IgG; R&D Systems). After rinsing, the sections were incubated with the following fluorescence-labeled secondary antibodies at room temperature for 1.5 h at a dilution of 1:500: Alexa Fluor 488-labeled donkey anti-rabbit IgG (A21206; Invitrogen), anti-rat IgG (A21208; Invitrogen), or anti-mouse IgG (A21202; Invitrogen). Images were captured under a FluoView FV10i confocal microscope (Olympus). The intensity of Iba1 immunoreactivity was measured across four fields of view (two lateral and two anterior funiculus of the spinal cord), each measuring 420 × 420 µm². The number of CD3-immunoreactive cells was calculated from the average number of cells in the right and left halves of the spinal cord. The number of Gr1-immunoreactive cells was counted in the anterior funiculus of the spinal cord within an 840 × 840 µm² field. Quantification was carried out using ImageJ software, as described previously [26].

### 2.5 Drug treatment

Based on its reported high oral bioavailability [27], AM095 was administered orally. For pre-onset EAE treatment, mice were allocated randomly to either the treatment or vehicle group on Day 3 post-immunization. Each mouse received 30 mg/kg AM095 (CS-1118; Chemscene) or vehicle (sterile water) on Days 3, 5, 7, 9, 11, and 13. For post-onset EAE treatment, mice were similarly assigned to groups on Day 16, and received 30 mg/kg AM095 or vehicle daily from Day 16 to Day 27.

### 2.6 Quantitative real-time PCR

Total RNA was extracted from the lumbar spinal cord (L3–L5) and spleen of both naïve and EAE mice using ISOGEN reagent (Wako), and cDNA synthesis was performed using ReverTra Ace® qPCR RT Master Mix (Toyobo). Real-time quantitative PCR was conducted with Power SYBR Green PCR Master Mix (Applied Biosystems) using the StepOne real-time PCR system (Applied Biosystems). The reactions were carried out in a total volume of 10 µL, which included 0.25 µg of total RNA and THUNDERBIRD SYBR qPCR Mix (Toyobo). The sequences of the oligonucleotide primers used are available in Supplementary Table 1.

### 2.7 Generation of bone marrow (BM) chimeric mice

C57BL/6-Tg(CAG-EGFP) mice (GFP^+^ mice) were purchased from Japan SLC (Shizuoka, Japan). LPA_1_-homo-KO mice were bred with C57BL/6-Tg(CAG-EGFP) mice to produce GFP^+^LPA_1_^+/−^ offspring. These GFP^+^LPA_1_^+/−^ mice were then crossed among themselves to generate GFP^+^LPA_1_^−/−^ mice. BM transplantation was performed as described previously, with minor modifications [26,28,29]. Briefly, female, 5-week-old C57BL/6J mice (WT) served as BM recipients. GFP^+^ mice or GFP^+^LPA_1_^−/−^ mice served as BM donors. Recipients were lethally irradiated for about 10 min with 8 Gy of whole-body irradiation. Donor mice were euthanized by decapitation and the femurs were separated carefully. Both ends of the femur were cut and BM cells were extracted using sterile PBS. The BM cell suspension was centrifuged at 400 × g for 10 min, and the resultant pellet of GFP^+^ BM cells was resuspended in sterile PBS. Within 3–5 h post-irradiation, WT recipient mice received an intravenous injection of 1.5 × 10^7^ BM cells into the tail vein. Two groups of WT recipient mice were established: those treated with GFP^+^ BM cells from GFP^+^ donor mice (WT^WT-BM^) and those receiving GFP^+^ BM cells from GFP^+^LPA_1_^-/-^ donor mice (WT^LPA1-KO-BM^). All mice were maintained under specific pathogen-free conditions, with unlimited access to autoclaved food and water. Six weeks later, these BM chimeric mice, now aged 11 weeks, were subjected to EAE induction.

### 2.8 Experimental design and statistical analysis

Statistical analysis was conducted using Prism 10 software (GraphPad Software). Details of the statistical methods and experimental design are provided in the legends and descriptions accompanying each figure. A mixed effects model with Šidák ’s post-hoc test was used to analyze EAE clinical scores, while one-way analysis of variance (ANOVA) with Dunnett’s post-hoc test was used to compare multiple experimental groups. Welch’s t-test was used to compare a single experimental group with a control group. *p* < 0.05 was considered statistically significant. Data are presented as the mean ± SEM. The evaluator was blinded to the treatment conditions. In the figures, each data point represents a sample (section or extract) from the lumber spinal cord or spleen of an individual mouse.

## 3. Results

### 3.1 Genetic deficiency of LPA_1_ reduces the pathological outcomes of EAE

To explore the role of LPA_1_ in MS, we immunized WT and LPA_1_-homo-KO mice [24] with a MOG 35-55 emulsion to induce EAE (Figure 1A). WT mice progressively developed severe EAE symptoms from the time of disease onset until the end of the experiment; by contrast, LPA_1_-homo-KO mice were resistant (Figure 1B). To further investigate involvement of LPA_1_ in EAE, we generated LPA_1_-hetero-KO mice and WT littermates. LPA_1_-hetero-KO mice tended to have lower clinical scores than their WT littermates (Figure 1C). To discern pathological differences between WT and LPA_1_-homo-KO mice, we employed immunohistochemical staining to assess the extent of demyelination and inflammatory cell infiltration in the CNS. In the spinal cord of LPA_1_-homo-KO mice, demyelination was notably reduced in the outermost white matter, as indicated by FluoroMyelin staining (Figure 1D and E). Additionally, fewer cells were observed in the spinal cord of these mice, as marked by DAPI staining (Figure 1D). These results suggest that the absence of LPA_1_ may mitigate demyelination and reduce the severity of EAE by limiting infiltration of peripheral immune cells, or by reducing activation of resident CNS cells.

**Figure 1.**
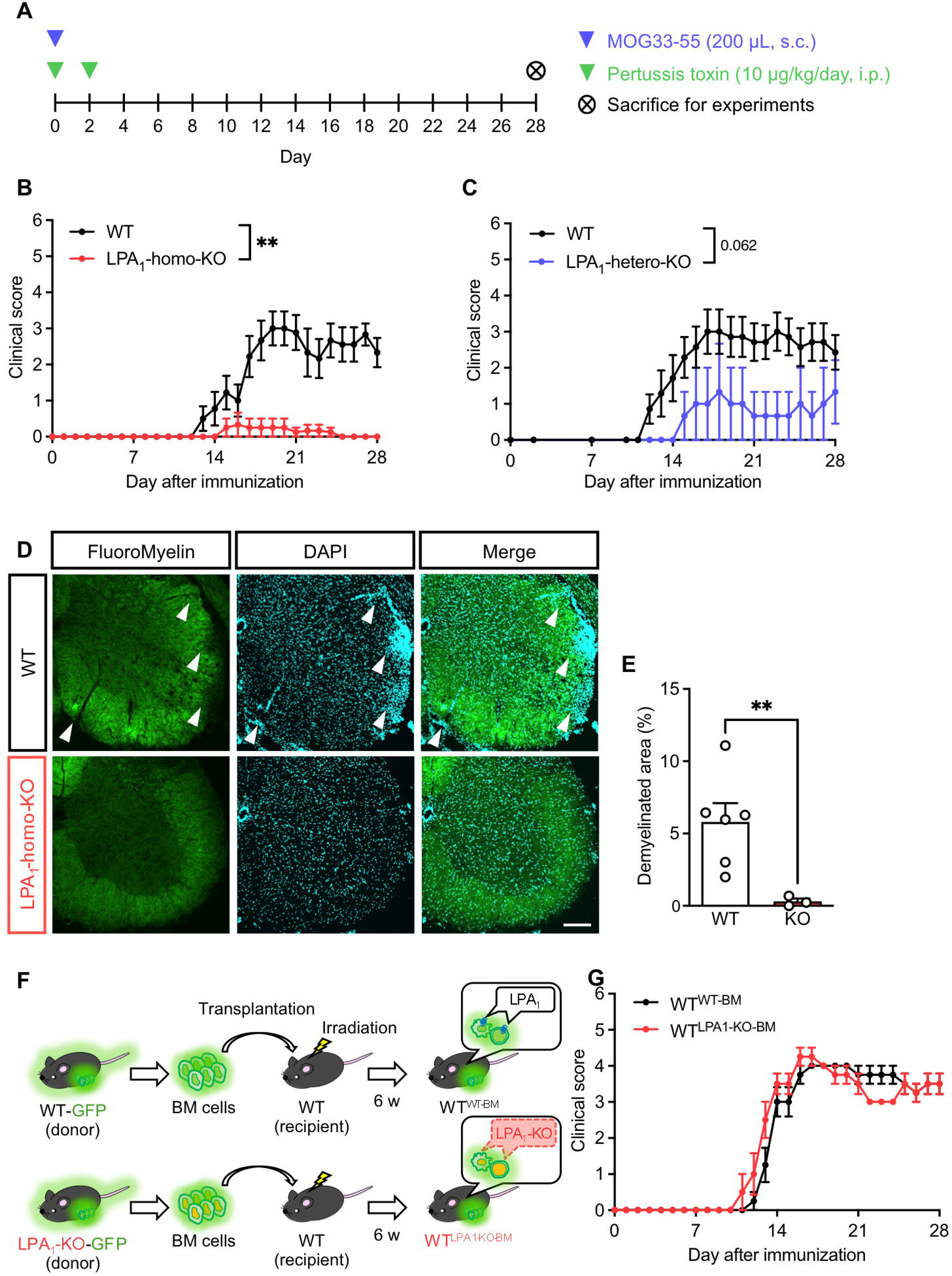
Genetic deficiency of LPA_1_ reduces the pathological outcomes of EAE. (A) Timeline of EAE induction. (B) Average clinical scores of LPA_1_-homo-KO and WT mice. *n* = 5–9. (C) Average clinical scores of LPA_1_-hetero-KO and WT mice. *n* = 3–7. (D) Representative images of lumbar spinal cord sections from WT and LPA_1_-homo-KO mice stained with FluoroMyelin to visualize demyelination and immune cell infiltration. White arrowheads point to areas of demyelination. Scale bar: 200 µm. (E) Quantitative analysis of the demyelinated areas depicted in (D). *n* = 3–6. (F) Schematic diagram illustrating the process used to generate BM transplant mice. (G) Average clinical scores of WT^WT-BM^ and WT^LAP1-KO-BM^ mice. *n* = 4. Mixed effects model with Šidák’s post-hoc (B, C, G); Welch’s *t*-test (E); \*\**p* < 0.01, \*\*\**p* < 0.001. Data are expressed as the mean ± S.E.M.

LPA_1_ is expressed in various tissues, including leukocytes such as dendritic cells [30] and macrophages [31,32]. These leukocytes play roles in the pathogenesis of EAE [26,33]. To investigate whether LPA_1_ expression in peripheral immune cells affects EAE pathogenesis, we generated BM chimeric mice by transplanting GFP^+^ BM cells from either GFP^+^ or GFP^+^LPA_1_^−/−^ donor mice into recipient mice that had undergone BM ablation through whole-body γ-ray irradiation (Figure 1F). Contrary to our expectations, the clinical scores of the WT^LPA1-KO-BM^ mice were similar to those of the WT^WT-BM^ mice (Figure 1G). The data suggest that LPA_1_ expression by peripheral immune cells does not contribute to EAE pathogenesis.

### 3.2 Genetic deficiency of LPA_1_ suppresses the increase in the number of microglia/macrophages in the spinal cord of EAE mice

The findings above indicate that LPA_1_ expression in CNS cells might contribute to the worsening pathology of EAE. Microglia/macrophages are involved in the exacerbation of MS pathology through axonal degeneration and myelin damage mediated by release of inflammatory cytokines and reactive oxygen and nitrogen species [34]. To investigate the impact of genetic deficiency of LPA_1_ on these cells, we stained the spinal cords of naïve WT, EAE-induced WT, and LPA**_1_**-homo-KO mice immunohistochemically. Immunoreactivity for Iba1, a marker for macrophages/microglia, was higher in EAE mice than in their naïve counterparts, but was reduced by genetic deficiency of LPA_1_ (Figure 2A and B). These results suggest that LPA_1_ plays a role in either activation of microglia, or in infiltration of the CNS by macrophages.

**Figure 2.**
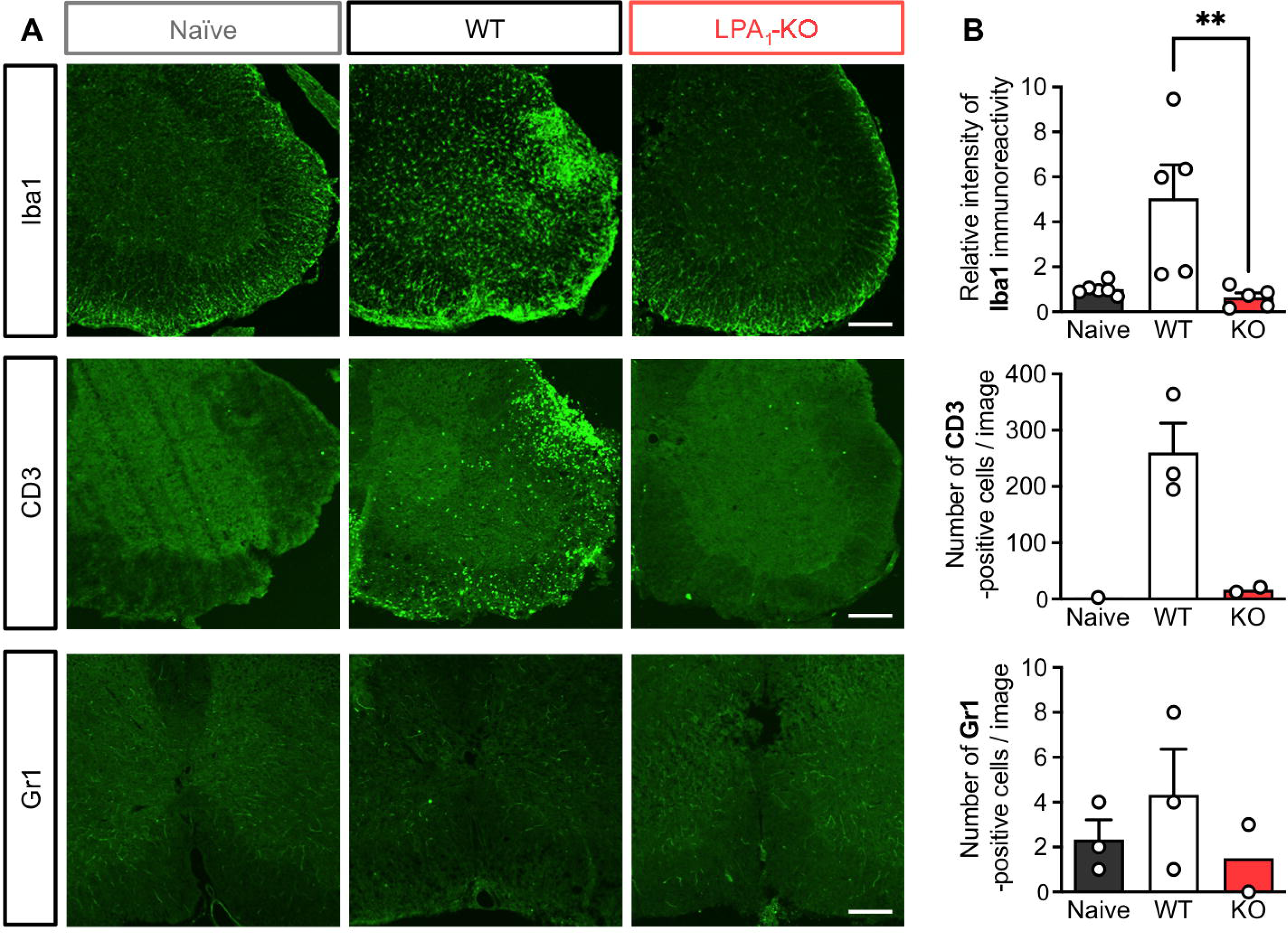
Genetic deficiency of LPA_1_ suppresses the increased numbers of microglia/macrophage in the spinal cord of EAE mice. (A) Representative images of lumbar spinal cord sections from naïve and EAE-induced WT and LPA_1_-homo-KO mice stained with anti-Iba1, anti-CD3, and anti-Gr1 antibodies. Scale bar: 200 µm for Iba1 and CD3; 100 µm for Gr1. (B) Quantitative analysis of the immunostained images displayed in (A). *n* = 8 for Iba1, *n* = 1–3 for CD3, *n* = 2–3 for Gr1. One-way analysis of variance (ANOVA) with Dunnett’s post-hoc test; \*\**p* < 0.01. Data are expressed as the mean ± S.E.M.

MS is associated with disruption of the blood-brain barrier (BBB), followed by infiltration by peripheral immune cells [35]. Th cells and neutrophils that infiltrate the CNS can either promote or protect against BBB breakdown by secreting various cytokines [35,36]. To assess the impact of LPA_1_ genetic deficiency on infiltration by peripheral immune cells, we analyzed expression of CD3, a T-cell marker, and Gr1, a neutrophil marker. The results showed a tendency toward a lower number of CD3-positive cells in EAE-induced LPA_1_-homo-KO mice than in their WT counterparts (Figure 2A and B); however, the number of Gr1-positive cells did not differ between EAE and LPA_1_-homo-KO mice (Figure 2A and B). These findings suggest that LPA_1_ may play a role in T-cell homing to the CNS during MS pathogenesis.

### 3.3 Administration of an LPA_1_ antagonist before EAE onset tends to confer resistance to EAE

To determine whether pharmacological inhibition of LPA_1_ affects progression of EAE, we administered the selective LPA_1_ antagonist AM095 (Figure 3A) [27,37] to WT mice. Administration of AM095 before EAE onset demonstrated a trend towards increased resistance to EAE compared with vehicle treatment (Figure 3B and C). Conversely, administration of AM095 after EAE onset did not reduce the severity of EAE when compared with vehicle treatment (Figure 3D and E). These findings indicate that LPA_1_ signaling plays a role in initiation, but not in progression, of EAE pathogenesis.

**Figure 3.**
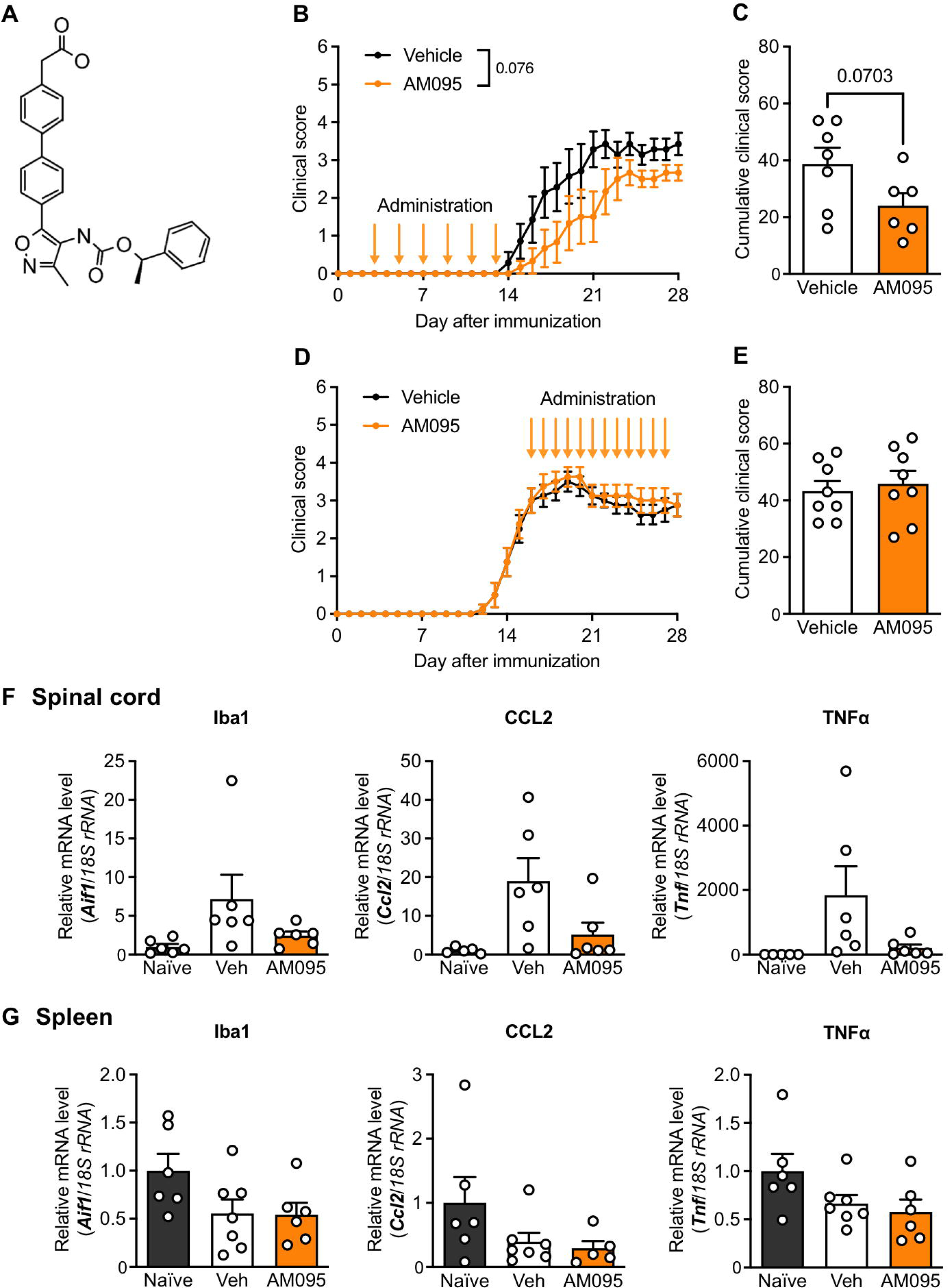
Administration of an LPA_1_ antagonist before onset of EAE tends to confer resistance to EAE. (A) Structural formula of AM095. (B, C) Average clinical score and cumulative clinical score of WT mice treated with AM095 or vehicle on Days 3, 5, 7, 9, 11, and 13. *n* = 6–7. (D, E) Average clinical score and cumulative clinical score of WT mice treated with AM095 or vehicle on Days 16–27. *n* = 8. (F, G) Molecular analysis post-treatment: Mice used in (B), as well as and naïve mice, were collected at Day 28 post-immunization and mRNA was extracted from the lumbar spinal cord and spleen. (F) Relative expression of mRNA in the lumbar spinal cord: *Aif1*, *Ccl2*, and *Tnf* (normalized against *18S rRNA*). (G) Relative expression of mRNA in the spleen: same markers as in (F). Each data point represents one marker per one sample per one mouse. Mixed effects model with Šidák’s post-hoc test (B, D); Welch’s *t*-test (C, E); One-way analysis of variance (ANOVA) with Dunnett’s post-hoc test (F, G). Data are expressed as the mean ± S.E.M.

Pharmacological inhibition of LPA_1_ after the onset of EAE did not ameliorate the disease, suggesting that LPA_1_ plays a crucial role in the pre-event phase of EAE. In MS patients, chemokines such as CCL2, CXCL8, and CXCL10 are involved in mobilizing peripheral immune cells [35], and cytokines such as IL1β, IL6, IL17, IFN-γ and TNFα contribute to the breakdown of BBB integrity [30]. Therefore, we asked whether administration of AM095 affects expression of these inflammatory cytokines and chemokines. Tissues from mice treated with AM095 before EAE onset were collected on Day 28, and mRNA expression levels in the spinal cord and spleen (key sites for immune cell maturation and storage) were examined. In the spleen, there was no change in the expression levels of mRNA encoding *Ccl2*, *Tnf*, *IL1b* and *Il6* in either the vehicle or AM095 groups (Figure 3G and 4D); however, in the spinal cord, expression of these mRNAs, which were elevated upon EAE induction, tended to decrease after treatment with AM095 (Figure 3F and 4C). This suggests that LPA_1_-mediated release of CCL2, TNFα, IL1β and IL6 may facilitate infiltration of peripheral immune cells and contribute to EAE pathogenesis.

**Figure 4.**
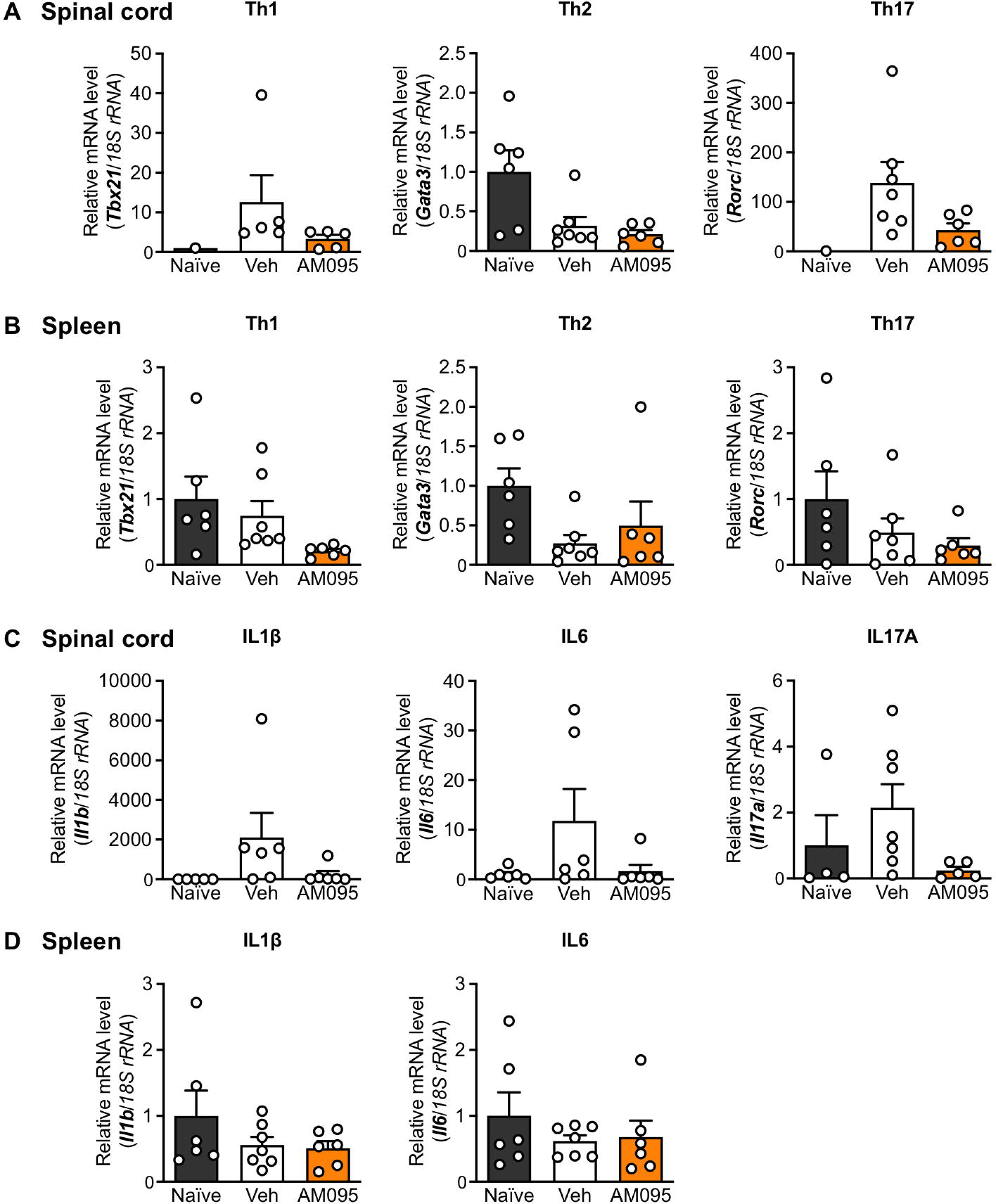
Pharmacological blockade of LPA_1_ reduces Th17 cell infiltration into the spinal cord of EAE mice. (A, B, C, D) Molecular analysis post-treatment. (A, C) Relative expression of mRNA in the lumbar spinal cord: *Tbx21*, *Gata3*, *Rorc*, *Il1b*, *Il6* and *Il17a* (normalized against *18S rRNA*). (B, D) Relative expression of mRNA in the spleen: same markers as in (A, C). Each data point represents one marker per one sample from each mouse. One-way analysis of variance (ANOVA) with Dunnett’s post-hoc test. Data are expressed as the mean ± S.E.M.

### 3.4 Pharmacological blockade of LPA_1_ reduces Th17 cell infiltration into the spinal cord of EAE mice

Suppression of central lymphocyte infiltration, a well-studied therapeutic target in the context of MS, focuses primarily on controlling the homing of B cells and T cells [38]. Among these lymphocytes, CD4-positive T cells, particularly Th1 and Th17 cells, play a crucial role in the mouse model of MS [39]. We investigated which helper T-cell subset among the CD3-positive T cells (which showed a trend toward reduced infiltration in Figure 2) was affected by AM095. Expression of mRNA encoding GATA3, a transcription factor specific for Th2 cells, and T-bet, a factor specific for Th1 cells, in both the spleen and spinal cord was no different between the vehicle and AM095 groups. Naïve T cells differentiate into Th17 cells in response to IL1β, IL6, IL23, and TNFα, and then secrete cytokines such as IL17A, IL17F, and IL21 [40,41]. Levels of *Il1b*, *Il6*, *Tnf* and *Il17a* mRNA (Figures 3F and G, and 4C and D) tended to decrease in spinal cords from the AM095 group compared with those from the vehicle group. Since differentiation of naïve CD4-positive T cells into Th17 cells typically does not occur in the CNS, this suggests that LPA_1_ does not affect differentiation of CD4-positive T cells; rather, it impacts infiltration of Th17 cells into the CNS and potentially increases secretion of IL17A.

## 4. Discussion

This study demonstrates that the pathogenesis of EAE is reduced significantly, and that demyelination is lessened, in mice lacking the LPA_1_ gene. LPA_1_-homo-KO mice exhibited less severe EAE than LPA_1_-hetero-KO mice, indicating that EAE severity may correlate with LPA_1_ expression levels. Conversely, expression of *Lpar1* mRNA decreases in the late stage of EAE group, a finding consistent with reanalyzed data from published single-cell RNA sequencing and single-nucleus RNA sequencing datasets [42,43] from both EAE mice and MS patients. This reduction in LPA_1_ expression might be a phenomenon manifested in the late stages of the disease as a result of a response by the biological defense system, although further research is necessary to clarify the underlying mechanisms.

Although conducted on a different genetic background, a previous report showed that the pathogenesis of EAE improves in the Malaga variant of LPA_1_-null mice [44]. That study also found that LPA_1_ expression by peripheral blood mononuclear cells (PBMCs) was elevated in a mouse model of MS, as well as in patients with relapsing-remitting MS, and that LPA stimulation leads to M1 polarization of macrophages [44]. The authors concluded that LPA_1_ expression in macrophages plays a role in the pathogenesis of EAE. Conversely, our results indicate that genetic deficiency of LPA_1_ in BM cells, including PBMCs and macrophages, does not ameliorate EAE pathogenesis (Figure 1G). The BM transplantation technique involves destruction of BM cells by whole-body γ-ray irradiation, yet the replacement of cells is limited predominantly to the periphery because resident CNS cells, including microglia, are resistant to myeloablative treatments [45]. These findings suggest that LPA_1_ expression in the CNS, rather than in peripheral immune cells, may make a more critical contribution to disease progression.

Genetic deficiency of LPA_1_ in EAE suppressed activation of Iba1-positive microglia in the spinal cord (Figure 2A and B). LPA_1_ is expressed by monocyte-derived cells, such as microglia and macrophages [31,32], In addition, LPA_1_ promotes microglial activation and TNFα production via the LPA_1_-ERK1/2 pathway [19]. Also, LPA_1_ expressed in microglia contributes to peripheral neuropathic pain [46,47]. Here, we found that *Tnf* mRNA levels tended to decrease in the spinal cord after AM095 administration. Additionally, the inflammatory mediators IL1β and IL6, as well as the chemokine CCL2 (which are commonly released from M1 microglia) [48,49] also showed a tendency to decrease at the mRNA level after AM095 administration. These findings suggest that LPA_1_ expressed presumably in microglia may contribute to the pathogenesis of EAE by promoting release of inflammatory mediators and chemokines.

Regarding LPA receptor subtypes, LPA_1_–LPA_6_ have been identified [18,50,51]. Specifically, genetic deficiency in LPA_2_ causes defects in lymphocyte homing, leading to severe exacerbation of EAE symptoms [30]. Conversely, agonists targeting LPA_2_ improve EAE symptoms [30]. Furthermore, studies indicate that administration of VPC32183 (an LPA_1_/LPA_3_ antagonist) improves EAE clinical symptoms [44], as does 1-oleoyl-LPA (an LPA_1_/LPA_2_ agonist) [52]. By contrast, Ki16425 (an LPA_1_/LPA_2_/LPA_3_ antagonist) exacerbates EAE symptoms [52]. These findings underscore the importance of understanding the specific roles of each LPA receptor subtype in the context of LPA signaling. LPA_1_ interacts with three G proteins: G_i/o_, which is involved in the RAS/MAPK and PI3K/AKT pathways, as well as in PLC activation; G_q/11_, which is also associated with PLC activation; and G_12/13_, which is linked to the Rho kinase pathway [53]. The transcription factor NF-κB, which acts downstream of the PI3K/AKT pathway, regulates production of CCL2, IL6, and TNFα. Therefore, microglia may release inflammatory mediators through the LPA_1_/PI3K/AKT pathway, thereby contributing to the inflammatory processes observed in diseases like EAE.

LPA production occurs via two primary pathways: direct hydrolysis of the fatty acid portion from membrane-derived phosphatidic acid, and ATX-mediated cleavage of the choline portion from lysophosphatidylcholine (LPC) [30]. ATX, an enzyme encoded by the *Enpp2* gene, is crucial for these processes, and its role in MS is actively being researched [11,12]. Genetic deficiency of ATX in CD11b-positive cells (microglia and macrophages) alleviates EAE pathology [12]. Upon activation, ATX interacts with cell surface integrins through its somatomedin B-like domains, thereby facilitating localized LPA production near target LPA receptors [54,55]. This mechanism allows LPA to act on receptors in a location-specific manner [30], suggesting that ATX deficiency in CD11b-positive cells (particularly microglia) may reduce LPA signaling in these cells, leading to decreased inflammation. Our re-analysis indicates that microglia express not only LPA_1_ but also LPA_5_ and LPA_6_ (Supplementary Figures 4–7). This diversity in receptor expression underscores the need for a more detailed exploration of the interactions between ATX and these LPA receptor subsets to fully understand their roles in inflammatory processes.

Genetic deficiency of LPA_1_ increased infiltration of the spinal cord by CD3-positive T cells. Corresponding with this observation, we found that administration of LPA_1_ antagonists before the onset of EAE tended to reduce elevated *Rorc* mRNA levels in the spinal cord, while not affecting Th17 cells in the spleen. LPA_1_ expressed on endothelial cells is implicated as a potential regulator of T-cell homing [56]. LPA_1_ may facilitate central homing of monocytes and T lymphocytes through mediators such as IL1β, IL6, and TNFα. Additionally, IL1β activates a variety of immune cells, including microglia and macrophages [57,58]. Once activated, microglia and macrophages contribute to demyelination and neuroaxonal degeneration through direct phagocytosis and release of inflammatory mediators [59,60]. Furthermore, IL17A, which is secreted by Th17 cells, plays a significant role in EAE pathogenesis [61–63]. LPA_1_ enhances the release of inflammatory mediators by activating immune cells within the CNS, thereby potentially exacerbating demyelination and neurodegeneration.

LPA_1_ is expressed by a variety of cells in different tissues, including the CNS and spleen [10,18,64]. While this study focused primarily on LPA_1_ in microglia/macrophages, LPA_1_ is also highly expressed in oligodendrocytes within the CNS [65–68]. This aligns with the findings from the re-analysis of published datasets [42,43,69,70] (Supplementary Figures 1-4). Genetic deficiency of LPA_1_ affects oligodendrocytes, leading to impaired cortical development, reduced levels of myelin basic protein (MBP), and thinning of the myelin sheath [71–73]. Recent research demonstrated that axonal insulation by oligodendrocytes is a risk factor for axonal degeneration, and that EAE clinical symptoms are unexpectedly less severe in mice with low MBP levels than in WT mice [74]. An additional hypothesis of this study suggests that loss of LPA_1_ results in decreased MBP levels, which may contribute to improvement in EAE pathology by preventing axonal degeneration. *Lpar1^flox/flox^* mice [47] and *Lpar1-EGFP* knock-in mice [75] may be valuable for determining which cells expressing LPA_1_ contribute to pathogenesis.

## DATA AVAILABILITY STATEMENT

The data that support the findings of this study are available from the corresponding author upon reasonable request.

## ACKNOWLEDGMENTS

This work was supported by Grants-in-Aid for Scientific Research (KAKENHI) from MEXT/JSPS (to H.S., JP23K27330, JP23H02639), and also by the Takeda Science Foundation and the Uehara memorial Foundation (to H.S.)

## Abbreviations

ATX: autotaxin
BM: bone marrow
CNS: central nervous system
EAE: experimental autoimmune encephalomyelitis
LPA: lysophosphatidic acid
MS: multiple sclerosis.

---

**Supplementary Table 1. Sequences of the oligonucleotide primers**

**Supplementary Figure 1. Re-analysis of published single-cell RNA sequencing data (Fournier AP et al., 2022** [39]**) derived from EAE mice.** The processed count data were retrieved from GSE199460 (available at NCBI GEO; https://www.ncbi.nlm.nih.gov/geo/query/acc.cgi?acc=GSE199460). The data were processed using Seurat v.5.0.3. Consistent with the methodologies described in the referenced paper, samples containing more than 25% mitochondrial genes, or those with fewer than 800 or more than 5500 features, were excluded from the analysis. Integration of each sample was performed using rpca (robust principal component analysis), focusing on the 2000 most variable features. Subsequently, nonlinear dimensionality reduction was executed using UMAP (Uniform Manifold Approximation and Projection), employing the first 20 principal components derived from PCA (principal component analysis). Clusters were annotated based on marker genes identified in the cited papers. (A) A violin plot displaying the expression level of the LPA_1_ gene (*Lpar1*) in each cluster. (B) A violin plot displaying the expression level of the ATX gene (*Enpp2*) in each cluster. (C) A dot plot visualizing the expression levels of LPA receptor genes (*Lpar1*, *Lpar2*, *Lpar3*, *Lpar4*, *Lpar5*, *Lpar6*) across the clusters.

**Supplementary Figure 2. Re-analysis of published single-nucleus RNA sequencing data (Jäkel S et al., 2019** [40]**) derived from patients with progressive MS.** The processed count data and metadata were retrieved from GSE118257 (available at the UCSC Cell Browser; https://cells.ucsc.edu/?ds=oligodendrocyte-ms). The data were processed using Seurat v.5.0.3. Consistent with the methodologies described in the referenced paper, samples containing more than 20% mitochondrial genes or those with fewer than 200 or more than 6000 features were excluded from the analysis. Integration of each sample was performed using CCA (Canonical Correlation Analysis) focusing on the 1000 most variable features. Subsequently, nonlinear dimensionality reduction was performed using UMAP, employing the first 30 principal components derived from PCA. Clusters were annotated based on metadata. (A) A violin plot displaying the expression level of the LPA_1_ gene (*LPAR1*) in each cluster. (B) A violin plot displaying the expression level of the ATX gene (*ENPP2*) in each cluster. (C) A dot plot visualizing expression levels of LPA receptor genes (*LPAR1*, *LPAR2*, *LPAR3*, *LPAR4*, *LPAR5*, *LPAR6*) across the clusters.

**Supplementary Figure 3. Re-analysis of published single-cell RNA sequencing data (Wheeler MA et al., 2020** [61]**) derived from EAE mice.** The processed count data were retrieved from GSE129609 (available at NCBI GEO; https://www.ncbi.nlm.nih.gov/geo/query/acc.cgi?acc=GSE129609). The data were processed using Seurat v.5.0.3. Three samples per group were used, and quality control was performed on each sample. Integration of each sample was performed using rpca, focusing on the 2000 most variable features. Subsequently, nonlinear dimensionality reduction was executed using UMAP, employing the first 20 principal components derived from PCA. Clusters were annotated using SingleR v.2.4.1, employing celldex MouseRNAseqData as the reference dataset. (A) A violin plot displaying expression of the LPA_1_ gene (*Lpar1*) in each cluster. (B) A violin plot displaying the expression level of the ATX gene (*Enpp2*) in each cluster. (C) A dot plot visualizing expression levels of LPA receptor genes (*Lpar1*, *Lpar2*, *Lpar3*, *Lpar4*, *Lpar5*, *Lpar6*) across the clusters.

**Supplementary Figure 4. Re-analysis of published single-cell RNA sequencing data (Falcão AM et al., 2018** [62]**) derived from oligodendrocyte lineage cells from EAE mice.** The processed count data and metadata were retrieved from GSE113973 (available at the UCSC Cell Browser; https://cells.ucsc.edu/?ds=oligo-lineage-ms). The data were processed using Seurat v.5.0.3. Consistent with the methodologies described in the referenced paper, samples containing more than 2% mitochondrial genes. or those with fewer than 2500 features, were excluded from the analysis. Integration of each sample was performed using rpca, focusing on the 2000 most variable features. Subsequently, nonlinear dimensionality reduction was executed using UMAP employing the first 30 principal components derived from PCA. Clusters were annotated based on metadata. (A) A violin plot displaying the expression level of the LPA_1_ gene (*Lpar1*) in control and EAE groups. (B) A violin plot displaying the expression level of the LPA_1_ gene (*Lpar1*) in each cluster. (C) A violin plot displaying the expression level of the ATX gene (*Enpp2*) in control and EAE groups. (D) A violin plot displaying the expression level of the ATX gene (*Enpp2*) in each cluster. (E) A dot plot visualizing the expression levels of LPA receptor genes (*Lpar1*, *Lpar2*, *Lpar3*, *Lpar4*, *Lpar5*, *Lpar6*) across the clusters.

