## Supplementary material for "Lysophosphatidic acid receptor 1 influences disease severity in a mouse model of multiple sclerosis": Table1&Suppl Fig. 1-4

| Gene | Forward primer | Reverse primer |
|---|---|---|
| *18S rRNA* | 5’-GCA ATT ATT CCC CAT GAA CG-3’ | 5’-GGC CTC ACT AAA CCA TCC AA-3’ |
| *Aif1* | 5’-TGA GGA GCC ATG AGC CAA AG-3’ | 5’-GCT TCA AGT TTG GAC GGC AG-3’ |
| *Ccl2* | 5’-AAC TCT CAC TGA AGC CAG CTC T-3’ | 5’-GTG GGG CGT TAA CTG CAT-3’ |
| *Foxp3* | 5’-CCT GGT TGT GAG AAG GTC TTC G-3’ | 5’-TGC TCC AGA GAC TGC ACC ACT T-3’ |
| *Gata3* | 5’-GGC AGA ACC GGC CCC TTA TC-3’ | 5’-TGG TCT GAC AGT TCG CGC AG-3’ |
| *Gfap* | 5’-GGC GAA GAA AAC CGC ATC AC-3’ | 5’-CCC GCA TCT CCA CAG TCT TT-3’ |
| *Il1b* | 5’-TGA GCA CCT TCT TTT CCT TCA-3’ | 5’-TTG TCT AAT GGG AAC GTC ACA C-3’ |
| *Il6* | 5’-GTG GCT AAG GAC CAA GAC CA-3’ | 5’-TAA CGC ACT AGG TTT GCC GA-3’ |
| *Il10* | 5’-GCT GCC TGC TCT TAC TGA CT-3’ | 5’-CTG GGA AGT GGG TGC AGT TA-3’ |
| *Il17a* | 5’-TCA GGG TCT TCA TTG CGG TG-3’ | 5’-TCT TTA ACT CCC TTG GCG CA-3’ |
| *Lpar1* | 5’-GAG GAT GTC TCG GCA TAG TTC TG-3’ | 5’-ATA AAG GCA CCA AGC ACA ATG A-3’ |
| *Tbx21* | 5’-CGG AGC GGA CCA ACA GCA TCG TTT C-3’ | 5’-CAG GGT AGC CAT CCA CGG GCG GGT-3’ |
| *Tgfb1* | 5’-TGA CGT CAC TGG AGT TGT ACG G-3’ | 5’-GGT TCA TGT CAT GGA TGG TGC-3’ |
| *Tnf* | 5’-TGC CTA TGT CTC AGC CTC TTC-3’ | 5’-GAG GCC ATT TGG GAA CTT CT-3’ |
| *Pdgfra* | 5’-GGG GAG AGT GAA GTG AGC TG-3’ | 5’-CAT CCG TCT GAG TGT GGT TG-3’ |
| *Rorc* | 5’-GAC AGG GAG CCA AGT TCT CA-3’ | 5’-CTT GTC CCC ACA GAT CTT GCA-3’ |

Uemura et al., Table 1

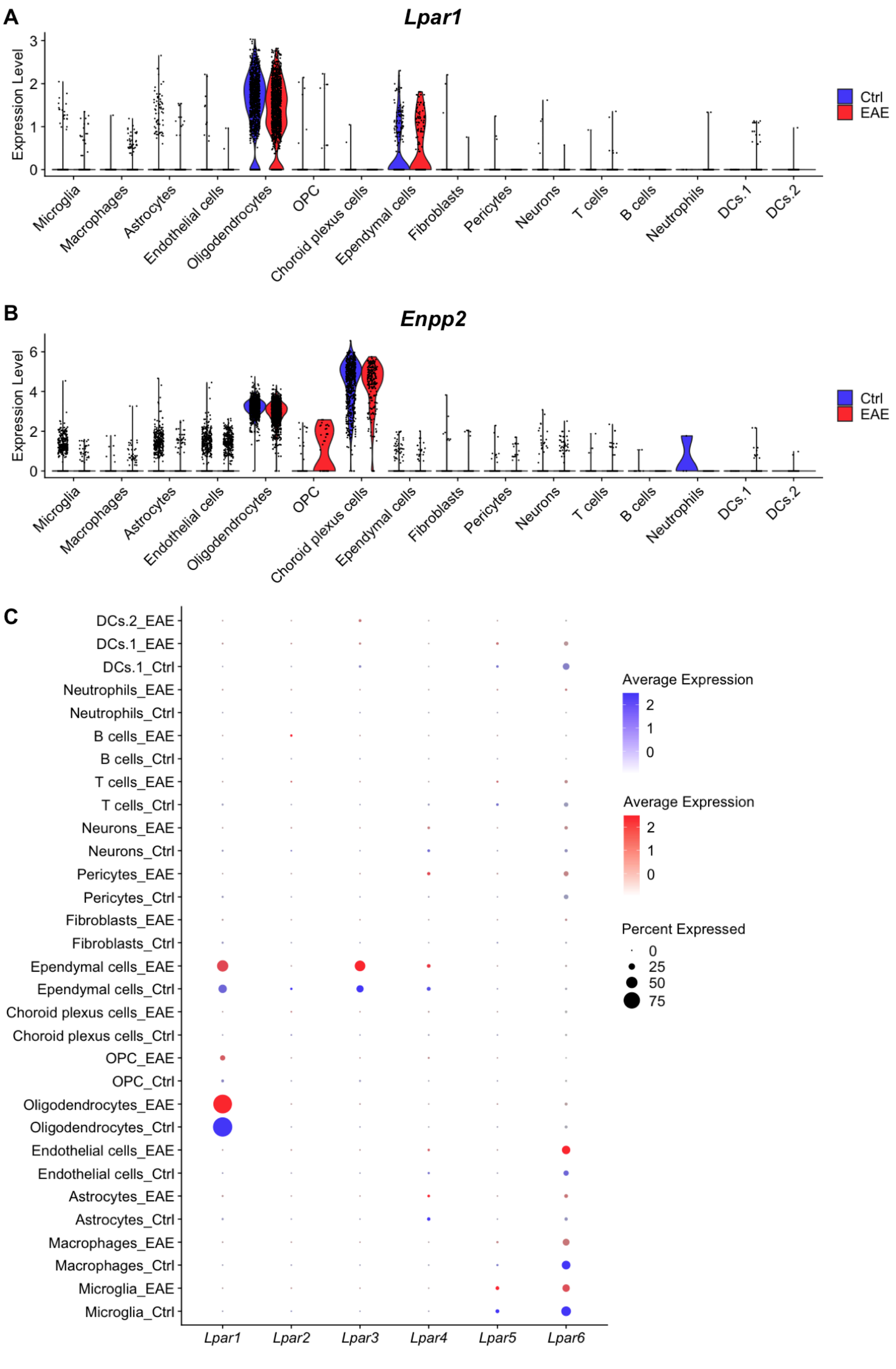


Uemura et al., Supplementary Figure 1

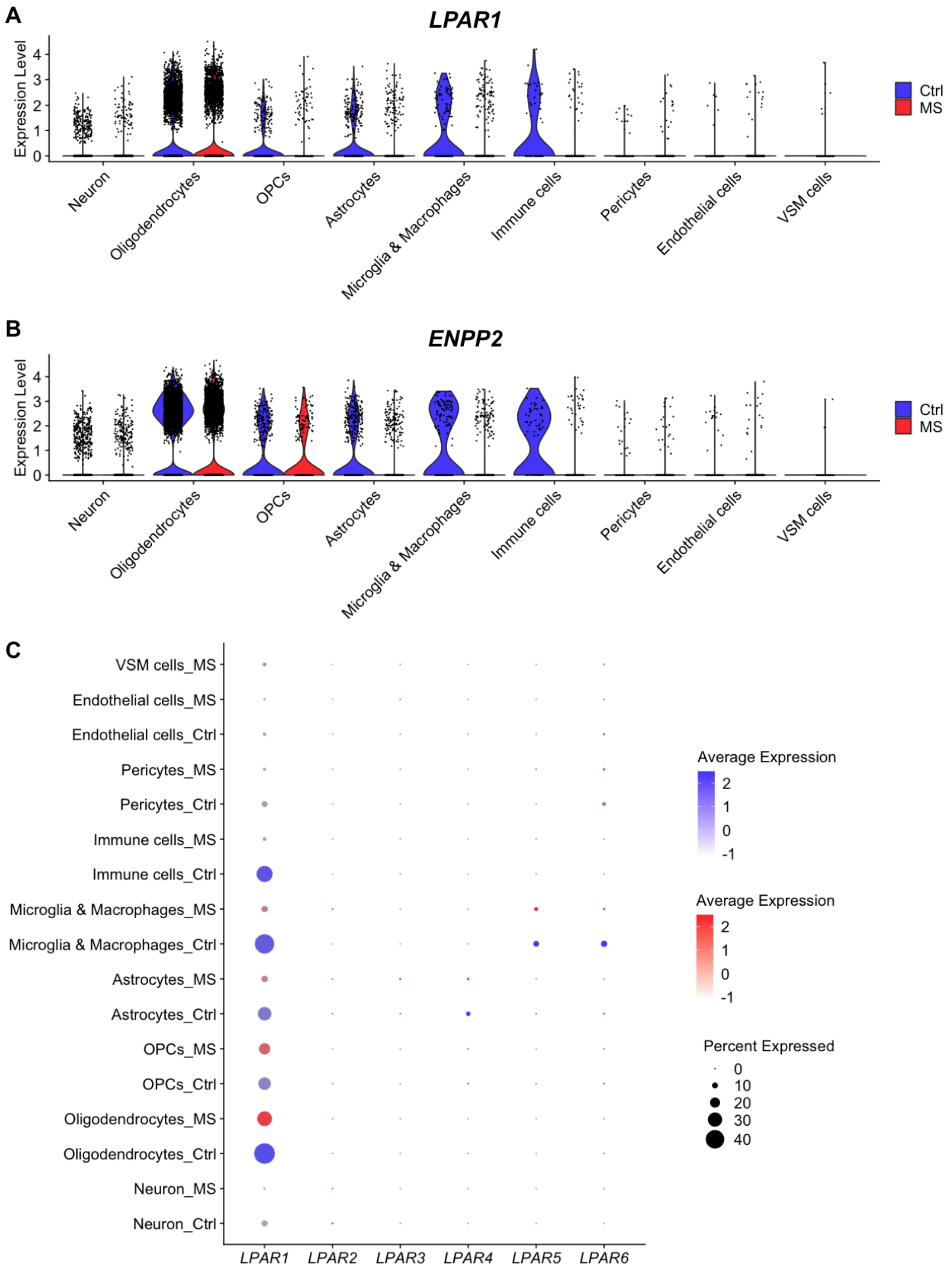


Uemura et al., Supplementary Figure 2

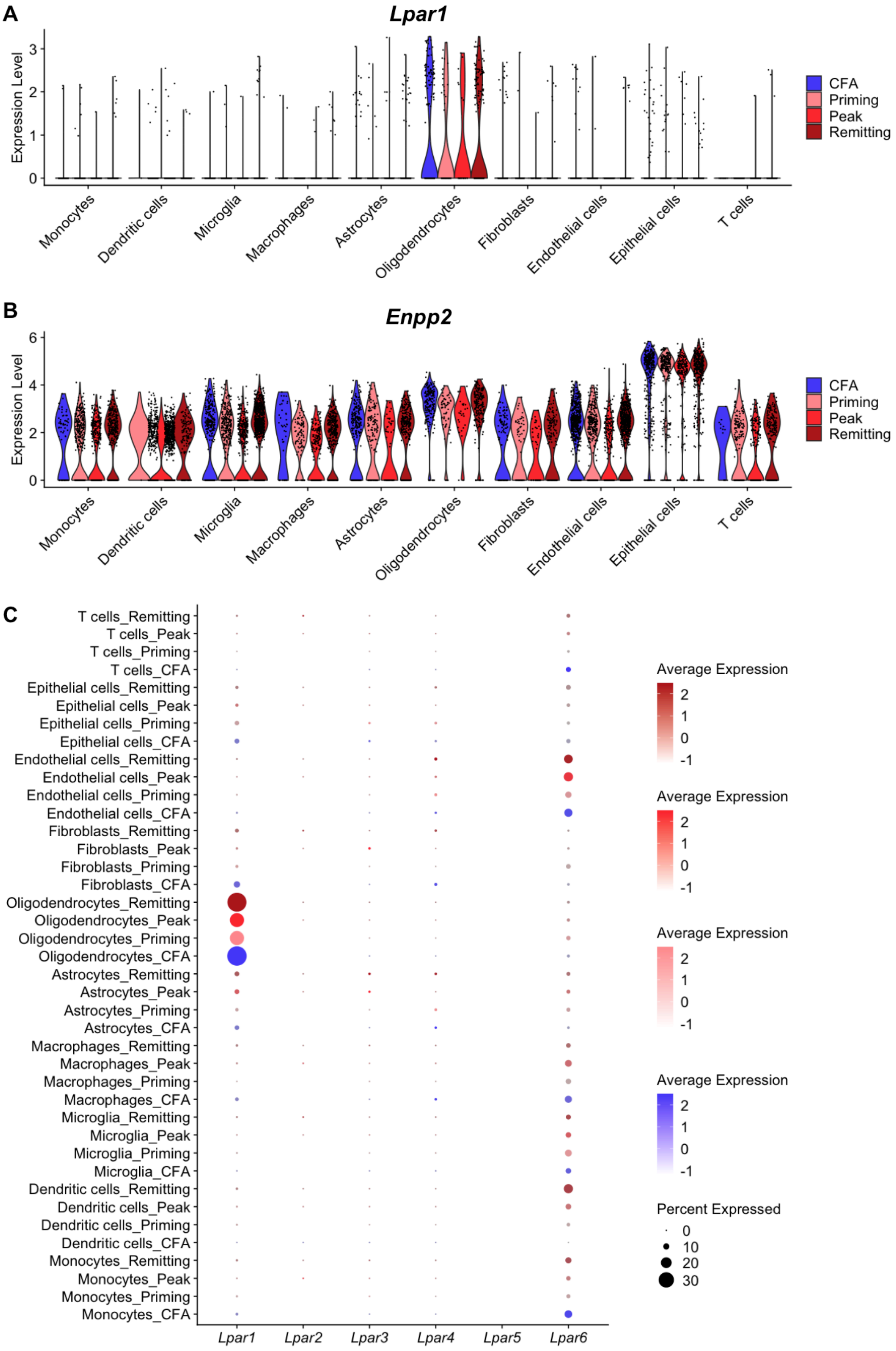


Uemura et al., Supplementary Figure 3

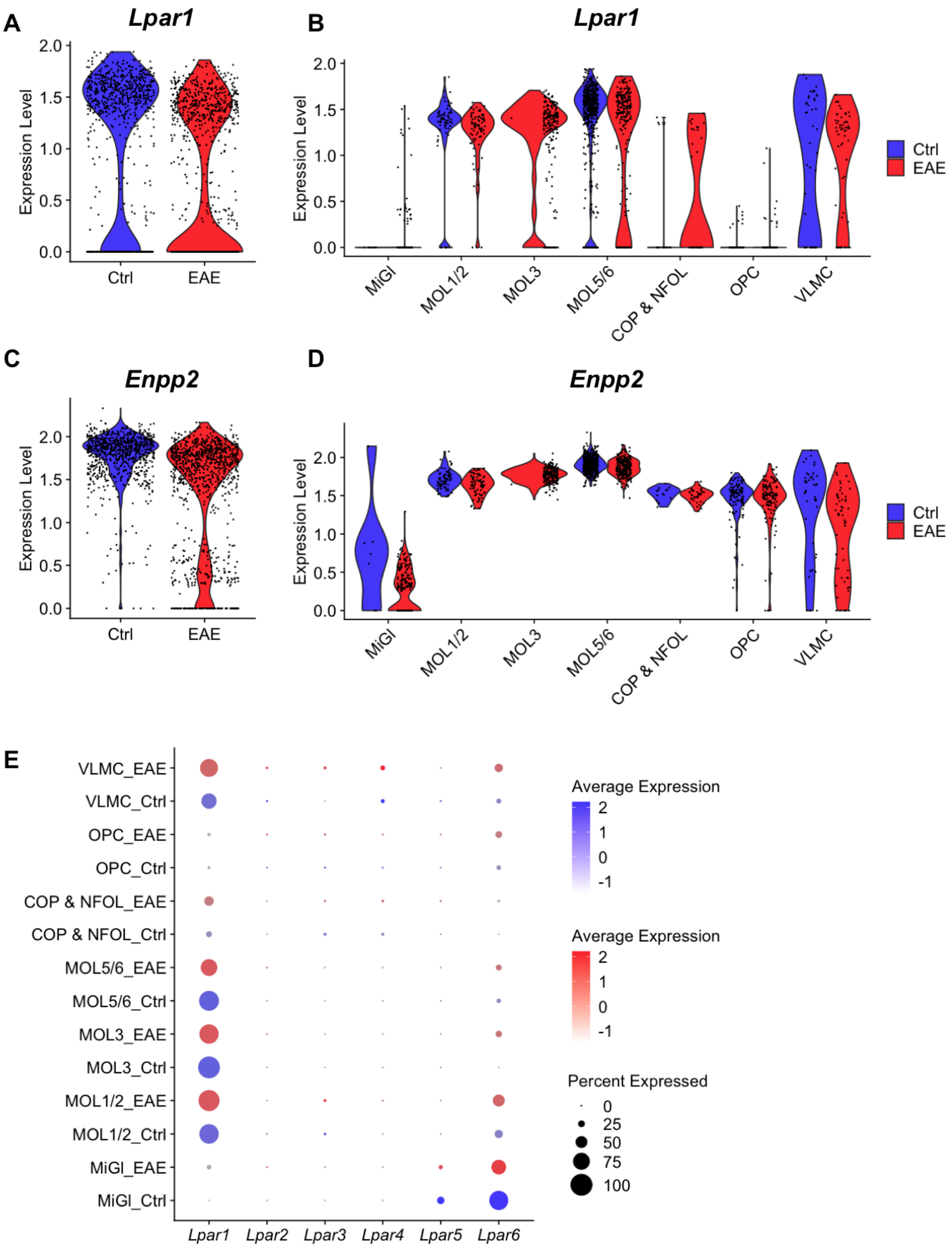


Uemura et al., Supplementary Figure 4
